# Dynamic Representation of the Subjective Value of Information

**DOI:** 10.1101/2021.02.12.431038

**Authors:** Kenji Kobayashi, Sangil Lee, Alexandre L. S. Filipowicz, Kara D. McGaughey, Joseph W. Kable, Matthew R. Nassar

## Abstract

To improve future decisions, people should seek information based on the value of information (*VOI*), which depends on the current evidence and the reward structure of the upcoming decision. When additional evidence is supplied, people should update *VOI* to adjust subsequent information seeking, but the neurocognitive mechanisms of this updating process remain unknown. We used a modified beads task to examine how the *VOI* is represented and updated in the human brain. We theoretically derived, and empirically verified, a normative prediction that the *VOI* depends on decision evidence and is biased by reward asymmetry. Using fMRI, we found that the subjective *VOI* is represented in right dorsolateral prefrontal cortex (DLPFC). Critically, this *VOI* representation was updated when additional evidence was supplied, showing that DLPFC dynamically tracks the up-to-date *VOI* over time. These results provide new insights into how humans adaptively seek information in the service of decision making.

## Introduction

Information seeking is critical for adaptive decision making. In order to improve future decisions, we collect information that would help us predict their outcomes. For instance, we check the weather forecast to decide whether to go out for a hike; we read about the policies and characters of candidates to decide how to vote; and we look up the number of COVID-19 cases to decide whether to have a family gathering. Recent work raises the possibility that deficits in information seeking underlie some psychiatric diseases such as schizophrenia and obsessive-compulsive disorder (OCD) (Baker et al., 2019; Dudley et al., 2016; Hauser et al., 2017; Ross et al., 2015).

In economic theories, information seeking should be primarily driven by information’s instrumentality, or how much the information would help the agent acquire rewards and avoid punishments in an upcoming decision. The information’s instrumentality is formally characterized as the value of information (*VOI*), defined as the improvement in the expected value (*EV*) that the agent can achieve by making the decision based on the information (Edwards, 1965; Howard, 1966). While this normative *VOI* theory does not incorporate psychological motives of curiosity, such as anticipatory utility (Caplin & Leahy, 2001; Gottlieb & Oudeyer, 2018; Kakade & Dayan, 2002; Kidd & Hayden, 2015; Kobayashi et al., 2019; Kreps & Porteus, 1978; Sharot & Sunstein, 2020), it predicts human participants’ information-seeking decisions reasonably well in settings where they acquire information at a cost (such as monetary costs or opportunity costs) that can be used to maximize rewards (Edwards & Slovic, 1965; Kobayashi & Hsu, 2019; Shanteau & Anderson, 1972; Wendt, 1969; Wilson et al., 2014). The idea that information seeking is driven by the *VOI* is further supported by electrophysiological and neuroimaging evidence that the *VOI* is encoded in reward-related regions (e.g., nucleus accumbens, ventromedial prefrontal cortex) as well as anterior cingulate cortex (ACC) and dorsolateral prefrontal cortex (DLPFC) (Blanchard et al., 2015; Bromberg-Martin & Hikosaka, 2009, 2011; Brydevall et al., 2018; Charpentier et al., 2018; Gruber et al., 2014; Jepma et al., 2012; Kaanders et al., 2020; Kang et al., 2009; Kobayashi & Hsu, 2019; Krebs et al., 2009; Lau et al., 2020; White et al., 2019).

The notion of the *VOI* based on the information’s instrumentality has two important implications. First, the *VOI* should not be determined by how much the information would contribute to the accuracy of prediction on the *state* of the world, but rather how much it would help the agent maximize *rewards*. Therefore, the *VOI* depends on the upcoming decision’s reward structure, or how rewarding or punishing possible outcomes are (for instance, the value of a weather forecast depends on how much the hiker prefers different weather conditions; those who don’t mind hiking in the rain or snow may not value the weather forecast as much as those who do). Second, the *VOI* depends on decision evidence that the agent already possesses prior to information seeking (Loewenstein, 1994). The *VOI* tends to be smaller when the agent already has more evidence, because they may already know what to do and additional information is less likely to influence it (e.g., a hiker may not need to check the weather forecast if they have been already informed by other hikers that it is going to snow). Thus, the agent needs to combine the available decision evidence with the reward structure to assess the *VOI* and seek information adaptively.

Crucially, when the decision evidence available to the agent changes, the agent should dynamically update the *VOI* based on the most recent evidence. Situations requiring such updating are ubiquitous in the real world, either because the environment gradually supplies evidence over time (e.g., a recent weather forecast is more accurate than an old one) or because the agent sequentially samples multiple pieces of information (the hiker can check multiple sources of weather forecasts). Despite its importance, to the best of our knowledge, no study has examined how the human brain tracks the up-to-date *VOI* based on the most recent decision evidence. The majority of neuroimaging studies so far have focused on cases where information is not instrumental for upcoming decisions, and those that have examined instrumentality-driven information seeking did not experimentally manipulate decision evidence over time to characterize the neural processes of updating the *VOI* (Kaanders et al., 2020; Kobayashi & Hsu, 2019).

We conducted an fMRI study to examine how human information-seeking behavior is sensitive to reward structure and current decision evidence, and how human brains track the up-to-date *VOI* after acquiring additional evidence. Our contributions are three-fold. First, we theoretically derive, and empirically demonstrate, a simple and generalizable prediction for how information seeking should be biased by asymmetry in reward structure. Second, we show that the right DLPFC represents the subjective *VOI* as a function of asymmetric rewards and current evidence. Third, we show that the *VOI* representation in the right DLPFC is dynamically updated when a new piece of evidence is supplied. These results suggest that the right DLPFC plays a critical role in information seeking in dynamic decision-making contexts by tracking the up-to-date *VOI* over time.

## Results

### Experimental paradigm

To examine neural representations of the value of information (*VOI*) and its updating, we adopted a variant of the beads task, an experimental paradigm widely used to study probability judgement and information seeking (Furl & Averbeck, 2011; Huq et al., 1988; Phillips & Edwards, 1966). As in the conventional version of the beads task, participants were presented with a jar containing two types of beads, one marked with a face and the other marked with a house, and asked to make a bet on its bead composition by observing some beads drawn from it. There were two possible compositions of the jar: one that consists of 60% face beads and 40% house beads, and the other that consists of 40% face beads and 60% house beads (Fig. 1A).

**Fig. 1.**
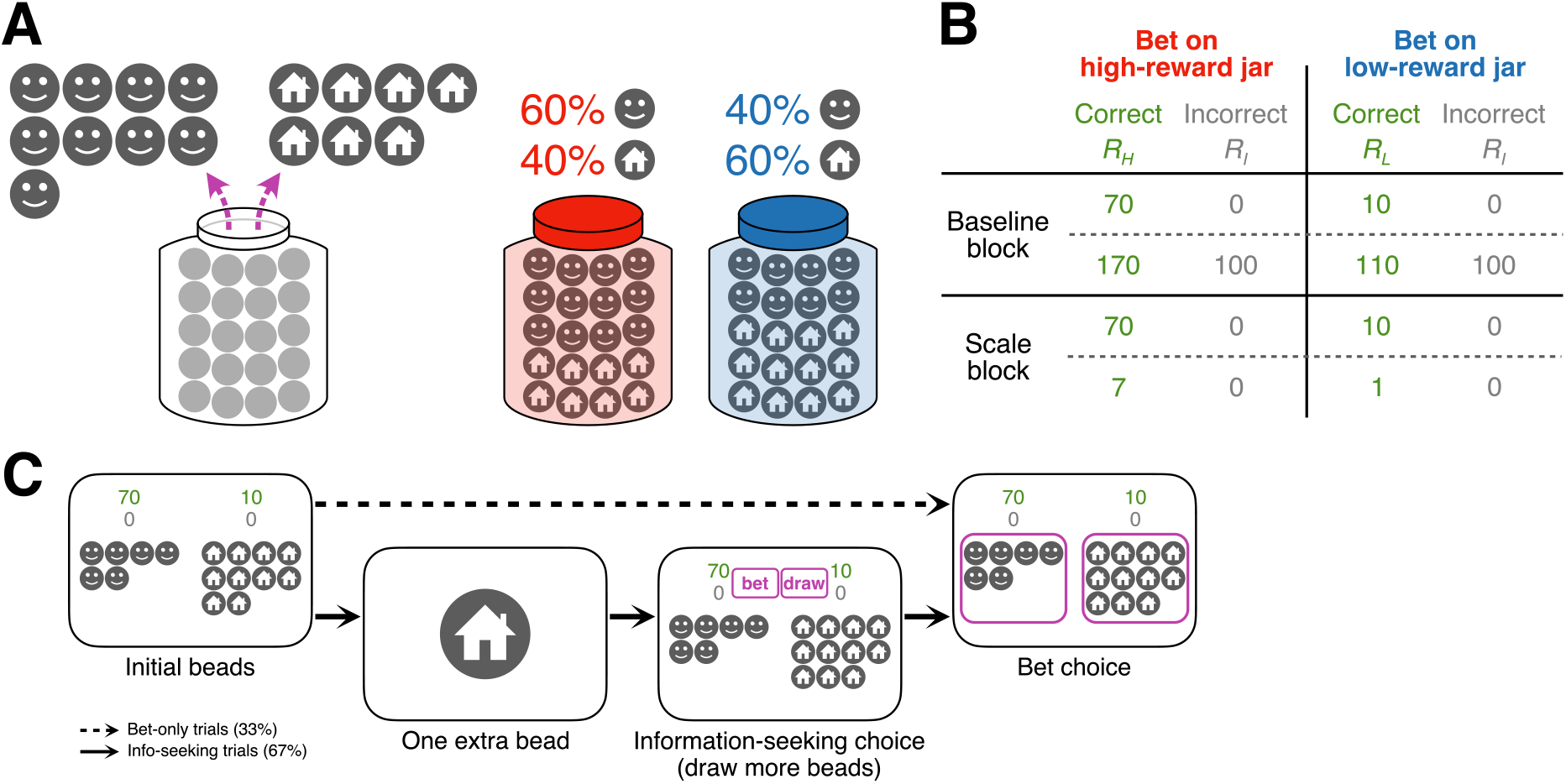
Experimental paradigm. We adopted the beads task with three key modifications: asymmetry in the reward structure, initial evidence prior to information seeking, and an updating event (one extra bead). (A) Participants observed a number of beads drawn from a jar and made a bet on its composition. Each bead was marked with a face or a house. There were two possible jar compositions: 60% face beads and 40% of house beads, or 40% face beads and 60% house beads. The jars are colored here only for illustrative purposes. (B) Reward structure. Participants earned more reward points by correctly betting on one of the two jar types. The experiment consisted of two blocks, in each of which one of the two reward structures was presented in each trial. The first block involved a baseline shift and the second block involved a scale manipulation. (C) Trial sequence. In a third of the trials (bet-only trials), participants were presented with a number of beads from the jar and immediately made a bet on its type. In the remaining trials (information-seeking trials), they were presented with the initial beads, an extra bead, and then allowed to seek further information by drawing more beads from the jar before making a bet on one of the two jars. Participants could draw as many beads as they needed within five seconds, but each additional draw incurred a cost (0.1 points). The extra bead was presented to evoke updating in the value of information.

Our variant of the beads task had three key features. First, we introduced reward asymmetry, such that participants could earn more reward by correctly betting on one jar type (e.g., the face-majority jar) than the other (e.g., the house-majority jar) (Fig. 1B). If participants were motivated to seek information to maximize rewards in the bet, their information-seeking strategy should be sensitive not only to the current evidence (the numbers of observed beads so far) but also to the reward asymmetry (the jar type they should bet on to maximize rewards). On the other hand, if participants were motivated to accurately guess the jar type, their information seeking should not be sensitive to the reward asymmetry. Therefore, the reward asymmetry allowed us to test whether information seeking was driven by the instrumentality of information for future reward seeking, as normatively prescribed in economic theories.

Second, we provided initial evidence, in the form of 20 or 30 bead draws from the jar. On a subset of trials, participants could then seek more information about the jar composition by drawing additional beads or elect to make a bet on the jar type (Fig. 1C). The difference in the numbers of face beads and house beads was parametrically manipulated to range from strong evidence favoring the low-reward jar to strong evidence favoring the high-reward jar. Additional draws incurred a small constant cost (0.1 points per draw) to monetarily incentivize participants to seek information only when necessary. This design allowed us to empirically measure the subjective *VOI,* or how much participants were willing to seek costly information, as a function of the current evidence.

Third, on the trials that allowed for information seeking, participants were presented with one extra bead draw from the jar prior to the information-seeking phase (Fig. 1C). The extra bead complemented the initial beads, shifting the evidence on the jar compositions, and thus updated the *VOI* originally evaluated based on the initial beads. We analyzed neural responses upon this extra bead event to examine how the neural representation of the *VOI* is dynamically updated based on the up-to-date evidence over time.

Participants completed the task inside the scanner. In each trial, after the presentation of initial beads and an extra bead, participants were allowed to draw as many additional beads as they wanted within five seconds, and then made a binary bet on the jar type. Additionally, to empirically elucidate participants’ reward-seeking behavior in a way that is not contaminated by information seeking, participants were asked to make a bet on the jar type without information seeking in a subset of trials (bet-only trials). Lastly, to explore how information seeking is sensitive to rewards, we introduced trial-wise manipulation of the reward structures. Specifically, participants earned a baseline reward of 100 points, irrespective of their bet, in half of the trials in one block (henceforth the baseline block), and they earned a tenth of the rewards in half of the trials in the other block (henceforth the scale block). Importantly, the reward of a correct bet was asymmetric across all trials and blocks (Fig. 1B).

### Theory

We first derived a theoretical prediction on how agents should seek information to optimize their bet and maximize rewards. We obtained theoretical *VOI* under the assumption that the agent aims to maximize the expected value (*EV*) of their decision, which they evaluate based on posterior probability of the jar type inferred in a perfectly Bayesian manner.

The posterior of the jar type is determined by the numbers of high-reward beads (the majority bead in the high-reward jar, e.g., face) and low-reward beads (the majority bead in the low-reward jar, e.g., house) observed from the jar so far (Fig. 2A). The more high-reward beads have been drawn, the more likely the jar is the high-reward jar, and vice versa. More specifically, the posterior is determined by the difference in the numbers of observed beads (high-reward beads minus low-reward beads) (Fig. 2B; Eq. 1). When more high-reward beads have been observed than low-reward beads (the beads difference > 0), the probability of the high-reward jar is higher than the probability of the low-reward jar, and it increases with the beads difference. Conversely, when more low-reward beads have been observed (the beads difference < 0), evidence favors the low-reward jar.

**Fig. 2.**
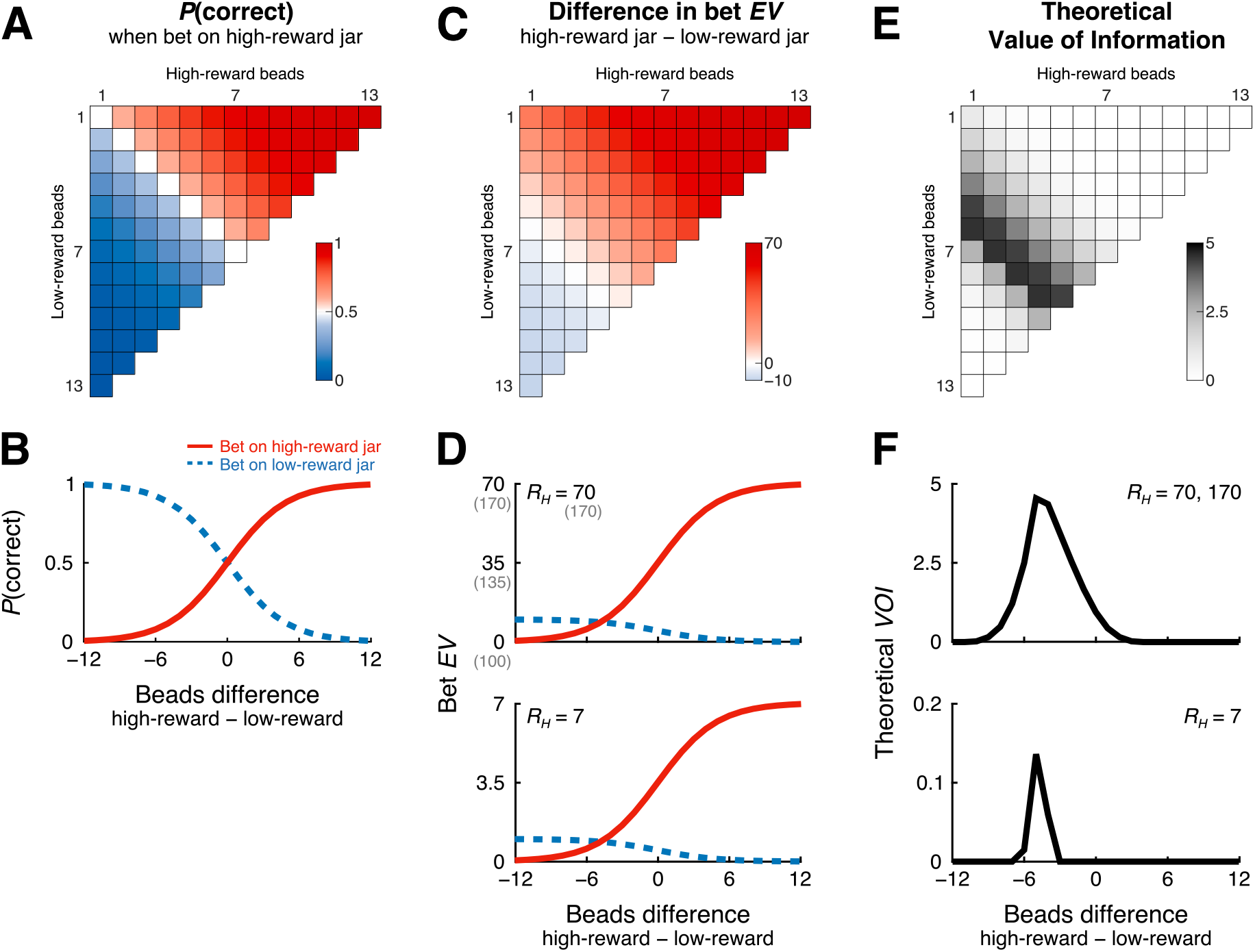
Theoretical predictions. (A) The probability of the jar type (the true jar is the high-reward jar) increases with the number of observed high-reward beads and decreases with the number of observed low-reward beads. (B) The probability of the jar type is determined by the beads difference. (C) Due to the reward asymmetry, when equal numbers of high-reward and low-reward beads have been observed (the diagonal), the *EV* to bet on the high-reward jar is higher than the low-reward jar. The agent would experience the smallest *EV* difference, and hence the highest uncertainty on the bet, when more low-reward beads have been drawn (the white region). (D) The *EV* difference is smallest at the beads difference of −5 across all reward structures. *Top*: Bet *EV*s are not affected by a baseline shift in rewards. *Bottom*: The relative magnitudes of *EV*s remain the same when rewards are scaled down overall. (E) The theoretical *VOI* is highest when the uncertainty on the bet is highest (the beads difference = −5, the black region). This is because the next bead would provide evidence in favor of either jar type, resolving the uncertainty. (F) The theoretical *VOI* takes an inverted-*U* shape across all reward structures. *Top*: The *VOI* is unaffected by a baseline shift in rewards. *Bottom*: When the rewards are scaled down, the magnitude of *VOI* becomes smaller as well, but the peak location remains the same. *EV*: expected value, *VOI*: value of information.

In order to evaluate the *EV* of a bet, the agent needs to combine the posterior on the jar type with the reward structure (Fig. 2C). Due to the reward asymmetry, when the current evidence does not favor either jar (the beads difference = 0; the diagonal in Fig. 2C), the *EV* to bet on the high-reward jar is higher than the *EV* to bet on the low-reward jar. The *EVs* to bet on the two jars are closest to each other when more low-reward beads have been observed (the beads difference = −5; the white region in Fig. 2C). This prediction holds across all of our reward structures (Fig. 2D); a baseline shift in rewards does not affect the *EV* difference, and a scale manipulation in rewards multiplicatively affect both *EV*s without changing their relative magnitudes. Therefore, if forced to bet on one of the two possible jars, the *EV*-maximizing agent would experience the highest choice uncertainty, not when equal numbers of beads have been observed, but when more low-reward beads have been observed than high-reward beads.

Under economic theories, the *VOI*, or the value of drawing an additional bead, is evaluated based on how much the next bead would improve the upcoming bet on average (Eq. 2). Qualitatively, the theoretical *VOI* tends to increase with the uncertainty about which jar type to bet on, because an additional bead would provide more evidence for either jar type and resolve the uncertainty over possible actions (Fig. 2E). For instance, when the agent is under high uncertainty on the bet (the beads difference = −5; the black region in Fig. 2E), an additional bead would help them make a bet *irrespective of its type*; if the next bead is a high-reward bead, it provides additional evidence in favor of the high-reward jar, whereas if it is a low-reward bead, it favors the low-reward jar. The agent can improve the *EV* by making a bet conditional on the next bead type. On the other hand, when the agent has observed more high-reward beads than low-reward beads (e.g., the beads difference = +10; top right in Fig. 2E), or when the agent has observed many more low-reward than high-reward beads (e.g., the beads difference = −10; bottom left in Fig. 2E), an additional bead would not affect the subsequent bet; the agent would bet on the high-reward jar or low-reward jar, respectively, no matter what the next bead would be. Therefore, the theoretical *VOI* takes an inverted-*U* shape as a function of the beads difference, with its peak at a negative beads difference (−5) (Fig. 2F).

Therefore, our theoretical framework yields an important prediction that the information-seeking strategy should be biased by the reward asymmetry; participants should draw additional beads more frequently when more low-reward beads have been observed than high-reward beads (the beads difference < 0). The predicted bias holds across reward structures (Fig. 2F); manipulation of the reward baseline (in the baseline block) does not affect the *VOI*, and manipulation of the reward scaling (in the scale block) affects the overall magnitude of the *VOI* but does not drastically alter its inverted-*U* shape. This prediction might be somewhat counterintuitive, as the motivation for information seeking is expected to be higher when the current evidence favors the less desirable state (the low-reward jar). However, it is consistent with the widespread notion of confirmation bias that an agent needs less evidence to bet on a desirable state than an undesirable state (e.g., Gesiarz et al., 2019). More generally, the prediction echoes the general assumption that information seeking should be driven not by the motivation to predict the *state* (which jar is the true jar?) but to maximize *rewards* (which jar to bet on?). If, in contrast to our theoretical assumption, an agent is solely motivated to accurately predict the state, they would seek information the most when the beads difference is zero. Therefore, a bias in information seeking would suggest that participants seek information based on its instrumentality for future reward seeking, as normatively prescribed. To our knowledge, the bias in information seeking under the reward asymmetry is a novel theoretical prediction that has not yet been directly tested.

### Behavior

We examined participants’ information-seeking behavior, and in particular, whether it was biased due to the reward asymmetry as predicted. If participants sought to improve their subsequent bet choice and maximize rewards, the frequency of information seeking (i.e., how often they drew at least one bead) should be biased towards a negative beads difference, i.e., when more low-reward beads have been drawn than high-reward beads.

Observed information-seeking behavior was biased in the predicted direction (Fig. 3A). In both baseline and scale blocks, the frequency of drawing an additional draw was highest when more low-reward beads had been drawn than high-reward beads. Sensitivity to the reward asymmetry was also confirmed by the bet on the jar type in the bet-only trials (Fig. 3B); the frequency of betting on the high-reward jar increased with the beads difference, and the indifference point (the point at which participants were equally likely to bet on either jar) was shifted towards a negative beads difference. These results show that participants incorporated both the current evidence and reward asymmetry in reward-seeking and information-seeking choices.

**Fig. 3.**
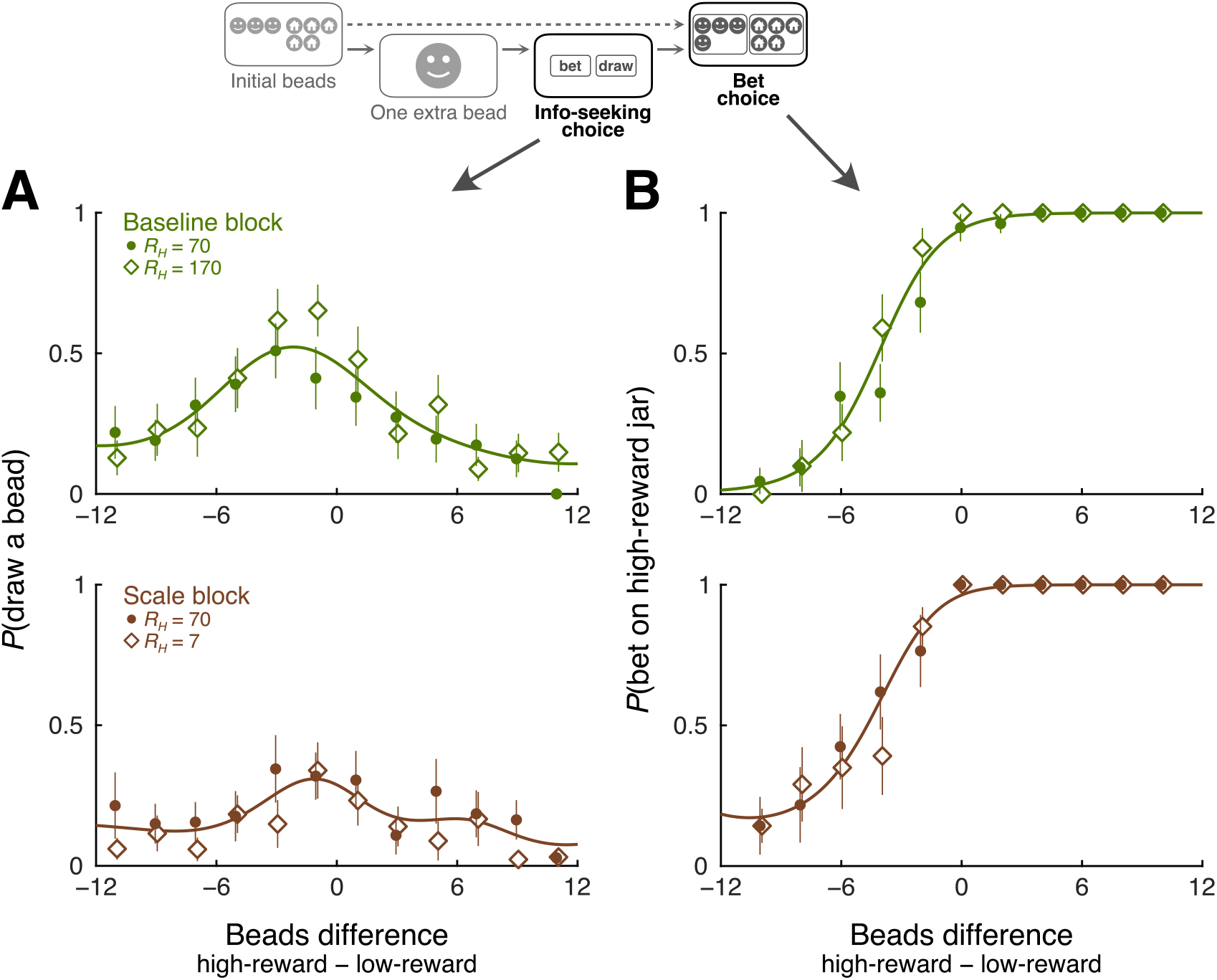
Behavior. Participants’ information-seeking and reward-seeking behavior was biased by the reward asymmetry as predicted. (A) Participants’ information seeking, or the frequency at which they drew at least one bead, peaked when more low-reward beads had been drawn than high-reward beads. (B) In the bet-only trials, the frequency with which they bet on the high-reward jar increased with the beads difference and was biased by the reward asymmetry. Lines indicate the best-fit model, which assumed sensitivity to blocks but not to reward manipulations within blocks. Error bars: bootstrap SD resampling participants.

A notable deviation from the theoretical prediction is that participants’ information seeking was not sensitive to the reward scale manipulation. In our framework, the theoretical *VOI* is smaller when the rewards are scaled down (even though its peak location remains the same) while it is unaffected by a reward baseline shift (Fig. 2F). Thus, if our participants were perfectly sensitive to the reward structure on a trial-by-trial basis, their information seeking should be affected by trial-wise reward manipulation in the scale block but not in the baseline block. To test this, we examined how information-seeking behavior differed across reward conditions and blocks. To characterize the relationship between information seeking and the beads difference without assuming its functional form, we used Gaussian Process (GP) logistic regression (Rasmussen & Williams, 2006). We fit four models to participants’ behavior; Model 1 assumed sensitivity to the scale manipulation but not to the baseline manipulation, as normatively prescribed; Model 2 assumed sensitivity to both manipulations; Model 3 assumed a difference between blocks but no sensitivity to manipulation in either block; and Model 4 assumed no difference between blocks or reward conditions. We found that Model 3 outperformed other models, including Model 1, according to both leave-one-participant-out cross validation (LOPO CV; log likelihoods [LL] = −1216.93, −1216.15, −1214.73, and −1232.15) and leave-one-trial-out cross validation (LOTO CV; LL = −1143.61, −1143.76, −1142.25, and −1166.17). Therefore, participant’s information-seeking behavior was systematically different between blocks, even though they did not change their strategy based on the reward structure on a trial-by-trial basis.

We speculate that shifting information-seeking strategies on a trial-by-trial basis was too cognitively taxing for our participants, because we also manipulated the beads difference and the trial type (information-seeking or bet-only). Despite this limitation, we observed that participants’ information seeking exhibited a clear bias in both blocks. Indeed, we observed that Model 3, which allowed asymmetry in information seeking, performed better than another model (Model 5) that assumed symmetric information seeking (baseline block LOPO CV LL = −666.82 [Model 3] vs. −679.52 [Model 5]; LOTO CV LL = −630.87 vs. −645.10; scale block LOPO CV LL = −547.92 vs. −548.13; LOTO CV LL = −511.68 vs. −512.53). Furthermore, analysis on betting choices also preferred Model 3 to Models 1 and 2 (comparison between Models 3 and 4 is equivocal; LOPO CV LL = −287.01, −285.67, −283.56, and −281.19; LOTO CV LL = −267.77, −266.25, −265.26, and −268.46), showing that participants were insensitive to trial-wise reward manipulation not only in information seeking but also in reward seeking. These results are qualitatively consistent with our theoretical prediction and lend support to the general notion that people seek information to improve their subsequent choices and maximize rewards.

### Neural representation of VOI

Next, we examined how the *VOI* was represented in the brain. Although previous fMRI studies reported *VOI* representations in a set of regions including DLPFC, VMPFC, and striatum, most of these studies focused on situations where participants obtained information that would not be useful for future decisions (i.e., information seeking for its own sake), and one study that examined instrumentality-driven information seeking used a one-shot paradigm that did not involve any updating (Kobayashi & Hsu, 2019). Thus, it remains unknown to what extent the neural representation of *VOI* is generalizable across tasks and decision contexts, and whether previously reported regions also represent and update the *VOI* in our experimental paradigm.

To look for brain regions that represent the *VOI*, we empirically estimated subjective *VOI* from the information-seeking behavior. We used the winning model of our GP logistic regression analysis (Model 3) to obtain the latent value function, which varied smoothly with the beads difference and differed between blocks (Fig. 4A). We then looked for regions where neural responses at the presentation of initial beads covaried with the subjective *VOI*.

**Fig. 4.**
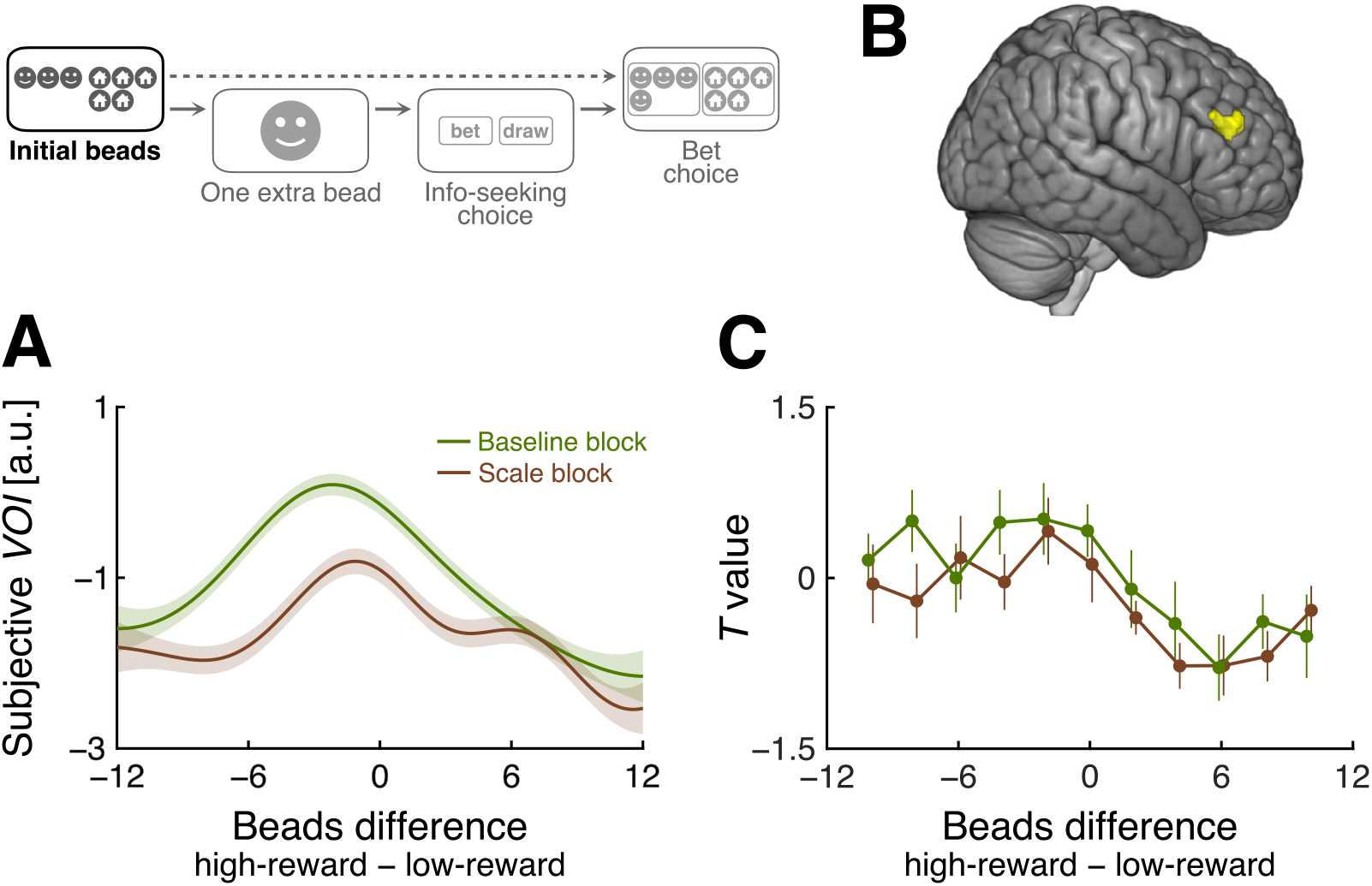
Neural representation of the *VOI*. (A) The subjective *VOI* was estimated for each block based on information-seeking behavior (Fig. 3A). (B) The right DLPFC represented the subjective *VOI* (cluster-mass *p* < .05, whole-brain FWE corrected; the peak MNI coordinate: [48, 42, 24]). (C) As predicted, the right DLPFC activation peaks at a negative beads difference in both blocks. Error bars: SEM.

We found a cluster in the right DLPFC representing subjective VOI (Fig. 4B; cluster-forming threshold *p* < .001, cluster-mass *p* < .05, whole-brain FWE corrected; peak MNI coordinate = [48, 42, 24]). Activation in this cluster peaked when more low-reward beads had been drawn in both blocks, consistent with the prediction (Fig. 4C). This cluster is the only one that survived our whole-brain statistical threshold (we also observed a cluster in the right anterior insula at a more lenient threshold, *p* < .10; peak MNI coordinate = [30, 24, 4]; Fig. S1).

Interestingly, the DLPFC cluster overlaps with a *VOI* cluster reported in a previous study that examined one-shot instrumentality-driven information seeking (Kobayashi & Hsu, 2019) (Fig. S1), providing converging evidence that the right DLPFC represents the *VOI* across decision contexts, at least when information is primarily acquired based on its instrumentality for future value-guided decisions.

### Updating of VOI representation

We then turned to our final question: how is the *VOI* updated upon the arrival of additional evidence in the brain? When the evidence available to agents changes, they need to track the up-to-date *VOI* in order to seek information adaptively over time. Specifically, we examined how the right DLPFC responds to the extra bead presented after the initial beads but prior to the information-seeking choice (Fig. 5A). We derived the *VOI* updating, or the difference between the posterior and prior *VOI*, as a function of the difference in the initial beads (the prior evidence) and the type of the extra bead (the evidence that causes updating). For instance, if participants have observed many more low-reward beads than high-reward beads (the beads difference < −5), an extra high-reward bead would positively update the *VOI*, as it slightly increases the uncertainty on the bet, while an extra low-reward bead would negatively update the *VOI*, as it further decreases the uncertainty on the bet. The directionality of updating is the opposite when more high-reward beads have been observed (the beads difference > 0).

**Fig. 5.**
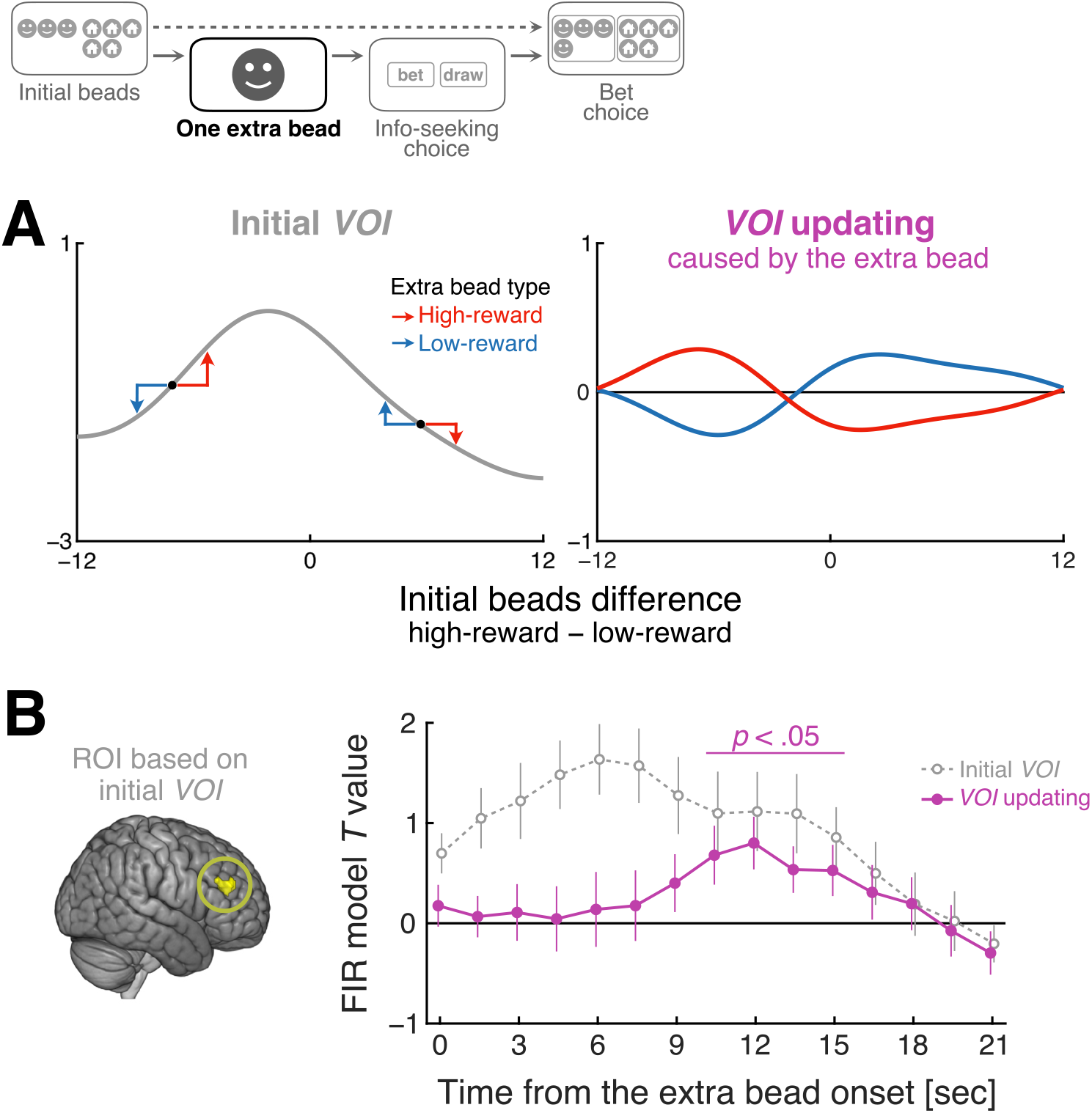
Updating of the *VOI* representation. The right DLPFC tracks *VOI* as it is updated by an extra bead, presented after the initial beads but prior to information seeking. (A) The *VOI* updating was calculated as the signed difference between the *VOI* after the extra bead and the *VOI* before the extra bead. (B) Time courses of the initial *VOI* signal (grey) and the *VOI* updating signal (purple) in the right DLPFC. The right DLPFC responds not only to the initial *VOI* but also to the updating of *VOI* (temporal cluster-mass *p* < .05, FWE corrected). Since the region of interest was defined based on the initial *VOI* signal, estimation of the initial *VOI* signal is biased, but estimation of the updating signal is unbiased. Error bars: SEM.

We hypothesized that the right DLPFC tracks the up-to-date *VOI* over time, such that it responds not only to the *VOI* based on the initial beads but is dynamically updated to the appropriate updated *VOI* after observation of the extra bead. To test this, we estimated the effects of the initial *VOI* and *VOI* updating on BOLD signals from the region of interest (ROI) defined above (Fig. 4B). In order to avoid a strong assumption about the time course of the updating process, we estimated the effects of initial *VOI* and *VOI* updating across time using finite impulse response (FIR) functions aligned to the presentation of the extra bead (Fig 5, top). We included three FIRs in a GLM, one parametrically modulated with the initial *VOI*, one modulated with the *VOI* updating, and one without parametric modulation (intercept). Since the ROI was originally defined based on its response to the initial *VOI* (albeit in an earlier time window), the estimated effect of the initial *VOI* is biased, but the estimated effect of the *VOI* updating depends critically on the exact bead that was drawn, and thus is independent of our ROI selection process (Fig 5A).

The estimated time courses are shown in Fig. 5B. As expected, the right DLPFC represents the initial *VOI* early on. Importantly, the right DLFPC also positively responded to the *VOI* updating (cluster-forming threshold *p* < .05, cluster-mass *p* < .05, FWE corrected across time). The rise of the *VOI* updating signal lags behind the initial *VOI* signal in time, but they go back to the baseline in parallel. The estimated time courses look somewhat sluggish, which presumably reflects the nature of our experimental paradigm in which participants had several seconds to complete information seeking.

This evidence demonstrates that neural representations in right DLPFC shift from the initial (*a priori*) *VOI* to the updated (*a posteriori*) *VOI*, suggesting that this brain region dynamically tracks the *VOI* based on the up-to-date evidence in service of adaptive information seeking over time.

## Discussion

In order to make better decisions, we need to seek information adaptively based on what we already know (up-to-date decision evidence) and what is at stake (reward structure). When our knowledge is updated, we need to update the *VOI* accordingly to decide whether to seek further information. Deficits in updating the *VOI* could lead to excessive repetition of information seeking even after enough evidence is accumulated (Hauser et al., 2017), or conversely, premature jumping to conclusions without enough evidence (Dudley et al., 2016; Ross et al., 2015). Despite its importance and ubiquity in the real world, we know little about how people evaluate and update the *VOI*. In this study, we used a variant of the beads task, in which decision evidence was parametrically manipulated on a trial-by-trial basis, to examine how information seeking is shaped by current evidence and asymmetric reward structure, and how the *VOI* is represented and updated in the brain.

We theoretically derived, and empirically verified, the normative prediction that information seeking should be biased by reward asymmetry. Participants were more likely to seek information when the current evidence preferred the less rewarding state due to high uncertainty on which state to bet. While the current study used asymmetric monetary rewards, our theoretical framework can be generalized beyond economic decision making based on the notion that the people assign intrinsic values to beliefs that they can hold (Kunda, 1991; Sharot & Garrett, 2016). If people are incentivized to hold certain beliefs, they will be more motivated to seek information when the current evidence supports the less desirable belief, even without extrinsic reward asymmetry (e.g., people check the latest number of COVID-19 cases more often when it is increasing than decreasing). It is worth noting, however, that the current study only examined reward structures where a correct bet yields asymmetric rewards but an incorrect bet does not, while outcomes of an incorrect prediction could also be asymmetric in some real-world scenarios (e.g., it would be more punishing to underestimate the chance of COVID-19 transmission and end up causing a superspreader event than to overestimate it and avoid a social gathering). More comprehensive, generalizable predictions would be obtained by expanding our findings to various reward structures.

Our theoretical and behavioral findings may provide some insight into confirmation biases observed across domains. Confirmation bias is commonly framed as biases in updating processes and/or decision criteria due to reward asymmetry or other factors such as pre-commitment (Gesiarz et al., 2019; Leong et al., 2019; Luu & Stocker, 2018; Talluri et al., 2018). We showed that, even without biases in updating or decision criteria, information seeking should be biased by reward asymmetry. The current study was not designed to test conventional confirmation bias; our behavioral measure of information seeking is not sensitive to a bias in updating, and a bias in decision criteria is not distinguishable from non-neutral risk attitude in our paradigm. Future research may examine how confirmation bias in updating and/or decision criteria affects information seeking, and conversely, how the information seeking bias would strengthen or weaken the effects of confirmation bias. Another exciting question for future research would be whether people exhibit an information-seeking bias upon sampling evidence from internal representations rather than the external world, such as episodic memory (Shadlen & Shohamy, 2016).

Our finding of the *VOI* representation in DLPFC is consistent with a previous fMRI study on instrumentality-driven information seeking (Kobayashi & Hsu, 2019), despite a number of key differences in task design. First, our paradigm required probabilistic inference on the hidden jar composition based on observable evidence, while Kobayashi & Hsu (2019) provided explicit and unambiguous visual presentation of outcome probability. Second, while Kobayashi & Hsu (2019) manipulated the information’s diagnosticity and cost on a trial-by-trial basis, the current paradigm did not (the participant always drew one bead at a time, which incurred a small constant cost). Third, and most importantly, unlike Kobayashi & Hsu (2019), the current study manipulated decision evidence available to the participant at the beginning of each trial and examined its effect on information-seeking behavior and underlying neural signals. Thus, the current study not only replicates but also critically extends Kobayashi & Hsu (2019)’s findings by showing that DLPFC is sensitive to the current evidence and biased by reward asymmetry, a key theoretical prediction of the instrumentality-driven *VOI*. Along with neuroimaging evidence that DLPFC is also activated upon information seeking driven by factors other than instrumentality (Gruber et al., 2014; Kang et al., 2009; Jepma et al., 2012), these results suggest that DLPFC is critical for adaptive information seeking across decision contexts and domains.

Unlike previous studies, we did not find *VOI* representation in reward regions (e.g., striatum or VMPFC) or ACC (Bromberg-Martin & Hikosaka, 2009, 2011; Brydevall et al, 2018; Charpentier et al., 2018; Gruber et al., 2014; Kaanders et al., 2020; Kang et al., 2009; Krebs et al., 2009; Lau et al., 2020; White et al., 2019). It is possible that we lacked statistical power to detect signals in these regions; indeed, we found a *VOI* cluster in anterior insula at a liberal threshold (Fig. S1), which often coactivates with ACC in task-based and resting-state fMRI (Fox et al., 2005; Menon & Uddin, 2010; Seeley et al., 2007). Alternatively, the involvement of these regions could depend on task and decision context. For instance, striatum and/or VMPFC may be more important when the information-seeking cost is larger and variable, which would demand online cost-benefit analysis (Lau et al., 2020; Kobayashi & Hsu, 2019). On the other hand, ACC may be more involved in evaluating uncertainty or conflict in the action space (Kennerley et al., 2011; Rudebeck et al., 2008; Rushworth & Behrens, 2008; Shenhav et al., 2016), which is tightly coupled with the *VOI* in many cases, particularly in situations that involve an exploration-exploitation tradeoff (Kaanders et al., 2020; Kolling et al., 2012; Shenhav et al., 2014). One possible reason that we did not observe representation of the *VOI* in ACC, at least at the standard statistical threshold we used, is that our experimental paradigm decoupled action uncertainty from the *VOI* computation in three ways: first, information-seeking trials were intermixed with bet-only trials and the participant could not tell the trial type upon the presentation of the initial evidence (the epoch where we observed the *VOI* representation; Fig. 4); second, the action uncertainty could not be evaluated until the presentation of the extra bead; and third, the information-seeking decision was mapped to different actions (left vs. right) across trials. Further research is needed to understand the extent to which functional localization of the *VOI* is dependent on task and decision context, and furthermore, how neural representation of the *VOI* is related to other forms of information seeking, including exploration and curiosity.

Importantly, we showed that DLPFC not only represents the *VOI* based on the initial evidence but also updates it when additional evidence is supplied, or in other words, DLPFC tracks the up-to-date *VOI* based on the most recent evidence. Such DLPFC signals may be critical for adaptive information seeking in situations where the agent accumulates decision evidence over time, either because it is gradually supplied from the environment or because the agent sequentially acquires multiple pieces of information. DLPFC may be well suited for sustained and dynamically updated representation of the *VOI*, as DLPFC neurons are known to exhibit sustained activity for working memory retention (Funahashi et al., 1989; Fuster & Alexander, 1971; Sreenivasan & D’Esposito, 2019). Critically, the *VOI* updating in DLPFC is distinct from information prediction error (IPE) signals observed in the dopaminergic reward system and habenula (Blanchard et al., 2015; Bromberg-Martin & Hikosaka, 2009, 2011; Charpantier et al., 2018); IPE encodes the probabilistic delivery of information itself, while the *VOI* updating is concerned with how the delivered information increases or decreases the instrumentality of further information. Exciting open questions for future research include whether *VOI* signals in DLPFC play a causal role in information-seeking behavior, and how they are adjusted when evidence acquired in the past becomes less relevant in a dynamic environment (Behrens et al., 2007; McGuire et al., 2014; Nassar et al., 2019).

Our results may have important implications for information-seeking deficits in clinical populations. For instance, schizophrenia has been associated with the tendency to make premature decisions without enough information seeking (Dudley et al., 2016; Ross et al., 2015; but see Baker et al., 2019), which could be accompanied by DLPFC hypoactivity (i.e., too low *VOI* signals) (Barch & Ceaser, 2012) and/or the lack of DLPFC’s sensitivity to current decision evidence and reward asymmetry. Similarly, OCD patients exhibit excessive information seeking (Hauser et al., 2017), which could be caused by hyperactivity in DLPFC (i.e., too high *VOI* signals) (Eng et al., 2015) and/or the lack of *VOI* updating in DLPFC. Our experimental and theoretical framework provides a novel approach to characterization of key components in instrumentality-driven information seeking, namely the sensitivity to current decision evidence, updating caused by additional evidence, and a bias due to reward asymmetry, which can be readily applied in future research with typical and clinical populations.

## Materials and Methods

All procedures were approved by the Institutional Review Board at the University of Pennsylvania.

### Participants

15 people (11 female, 4 male, age: 18-28, mean = 21.27, standard deviation = 2.79) participated in the experiment. They provided informed consent in accordance with the Declaration of Helsinki.

### Task design

We adopted a variant of the beads task (Furl & Averbeck, 2011; Huq et al., 1988; Phillips & Edwards, 1966); the participant was presented with a jar containing two types of beads and asked to guess its composition (i.e., which type made up the majority of the beads) by drawing some beads from the jar (Fig. 1A). Our variant had three important features. First, the participant was rewarded for identifying the correct jar composition, but the reward structure was asymmetric, such that the participant could earn more rewards by correctly betting on one jar type than the other (Fig. 1B). Second, a variable number of beads was drawn from the jar and presented to the participant at the beginning of each trial, empirically manipulating the evidence available to the participant before they seek information. Third, an extra bead was presented on a subset of trials to update the initial evidence. These features allowed us to examine how the brain represents and updates the *VOI* based on evidence that changes over time.

The experiment consisted of two interleaved trial types, bet-only trials and information-seeking trials (Fig. 1C). In the bet-only trials, the participant was first presented with a number of beads drawn from the jar. Each bead was depicted as a rounded picture of a face or a house (one picture for face or house each was used throughout the experiment). Beads marked with a face were presented to the left and those marked with a house to the right. The participant was told that these beads were drawn from one of two jars: a face-majority jar, which consisted of 60% face beads and 40% house beads, and a house-majority jar, which consisted of 60% house beads and 40% face beads. Rewards for correct and incorrect bets (in points) were also presented, in green and gray, respectively. Rewards for a bet on the face-majority jar were shown above the face beads, and rewards for a bet on the house-majority jar above the house beads. Rewards for a correct bet on one jar were numerically larger than rewards for a correct bet on the other jar (reward asymmetry), while an incorrect bet on either jar yielded the same rewards (Fig. 1B). After the presentation of the initial beads for 3 seconds, the participant was asked to make a bet. During the bet phase of the task, face and house beads were separately outlined by magenta boxes, and the participant could press the left or right button on a response box to bet on the face- or house-majority jar, respectively. Trials in which the participant did not make a bet within 3 seconds were terminated and discarded from the analysis.

In the information-seeking trials, the participant was first presented with the initial beads screen (same as the bet-only trials), followed by a blank screen (0-2 seconds). Next, an extra bead drawn from the jar was presented, either marked with a face or a house (1 second), which was added to the corresponding group of beads on the initial screen (0-2 seconds). The participant was then asked to decide whether to draw more beads from the jar before making a bet on its composition (information-seeking phase). Two choices appeared on the screen, “draw” and “bet”, and the participant pressed one button to draw one more bead and another button to terminate the information-seeking phase and proceed to the bet (the sides of the options were randomized across trials). The participant was allowed to draw as many beads as they wanted within 5 seconds, and a face or house bead was added to the screen every time they pressed the “draw” button. The participant was told that each draw incurred a constant small cost (0.1 points). Once they pressed the “bet” button (or when 5 seconds have passed), they were presented with the bet screen (same as the bet-only trials).

The task was programmed in Matlab (The MathWorks, Natick, MA) using MGL (http://justingardner.net/mgl/) and SnowDots (http://code.google.com/p/snow-dots/) extensions.

### Procedure

In a separate task session before scanning, participants received extensive training on the task, in which various aspects of the task were gradually introduced (betting on the jar composition, asymmetric rewards, costly draws, and multiple reward structures). During the subsequent session, participants completed the task inside the scanner. Participants made responses using an MRI-compatible button box. They were compensated based on the total points they acquired in the scanning session (500 points = $1).

The scanning experiment consisted of two blocks, which differed in reward structure (Fig. 1B). In the first block (the baseline block), one of the two reward structures, (*R_H_*, *R_L_*, *R_I_*) = (70, 10, 0) or (170, 110, 100), was randomly presented in each trial, where *R_H_* is the reward for a correct bet on the high-reward jar, *R_L_* is the reward for a correct bet on the low-reward jar, and *R_I_* is the reward for an incorrect bet; thus, the participant earned a baseline reward of 100 points irrespective of their bet in half of the trials. In the second block (the scale block), one of the two reward structures, (*R_H_*, *R_L_*, *R_I_*) = (70, 10, 0) or (7, 1, 0), was randomly presented in each trial; thus, the participant earned a tenth of the rewards in half of the trials. Each block consisted of two scanning runs, one where the high-reward jar was the face-majority jar and one where the high-reward jar was the house-majority jar; their order was counterbalanced across participants.

On each trial, the participant was presented with 20 or 30 initial beads from the jar. The difference in the number of initial beads marked with a face or house was uniformly sampled from a discrete set of values ranging from −10 to 10 in increments of 2. Unbeknownst to the participant, the true jar type was stochastically determined following the Bayesian posterior conditional on the initial beads difference (see Eq. 1 below). In the information-seeking trials, the type of the extra bead presented and of all additional beads drawn by the participant (face or house) were stochastically determined based on the hidden jar type. The participant was not provided with feedback on their bet accuracy or rewards on a trial-by-trial basis. They were however informed of the total number of points they had accumulated at the end of each run.

### Theory

Normative predictions about the *VOI*, or how much an optimal agent should pay for information, were derived under assumptions that the agent conducts full-Bayesian inference on the jar type, deterministically makes an optimal choice to maximize the expected value (*EV*), is risk neutral, and optimally seeks information based on its instrumentality, or how much it would improve the *EV* of the subsequent bet choice. Our theoretical framework did not consider any additional information-seeking motives, such as curiosity, savoring, dread, or uncertainty reduction.

Let *SH* be the state where the true jar is the high-reward jar and *SL* the state where it is the low-reward jar. Let *α_H_* be the action to bet on *S_H_* and *α_L_* the action to bet on *SL*. Let us further refer to the majority beads in the high-reward jar as high-reward beads and the majority beads in the low-reward jar as low-reward beads (for instance, if the high-reward jar is the house-majority jar, a house bead is a high-reward bead and a face bead is a low-reward bead; note that the beads were not directly associated with rewards per se). The goal for the agent is to choose between *α_H_* and *α_L_* to maximize *EV* given the current evidence (i.e., the number of high-reward beads *n_H_* and low-reward beads *n_L_* drawn from the jar so far) and the reward structure (*R_H_*, *R_L_*, *R_I_*).

The likelihood of drawing a high-reward bead *b_H_* or a low-reward bead *b_L_* conditional on the jar type is known to the agent:

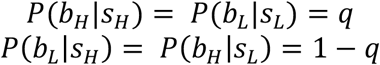

where *q* = 0.6. Assuming that the agent has a flat prior on the jar type (*P*(*S_H_*) = *P*(*S_L_*) = 0.5), the posterior follows

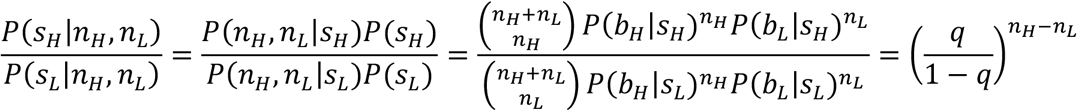

therefore

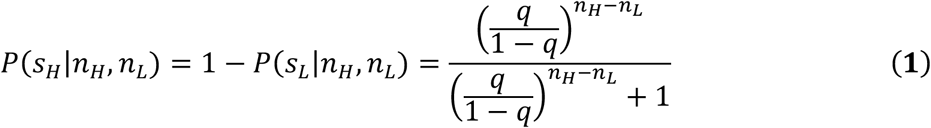

which is a function of the beads difference, *n_H_* − *n_L_* (e.g., the posterior is the same when (*n_H_*, *n_L_*) = (5, 2) or (15, 12)) (Fig. 2a, b).

Given the posterior, the agent makes a choice among three options: to bet on *S_H_*, to bet on *S_L_*, or to seek information and draw an additional bead from the jar, which incurs a cost *c*_draw_ (0.1 points). The agent should decide whether to draw an additional bead based on the *VOI*, or the improvement in the bet’s *EV* thanks to the next bead:

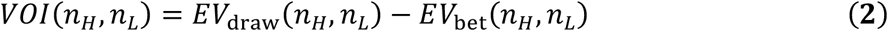

where *EV*_draw_ is the highest *EV* that the agent could achieve after drawing the next bead (without considering the information-seeking cost), and *EV*_bet_ is the highest *EV* that the agent could achieve by making a bet without any further information. The agent should draw a bead if and only if the *VOI* is higher than the drawing cost *c*_draw_.

*EV*_bet_ is the higher of the two bet *EV*s based on the current evidence, namely

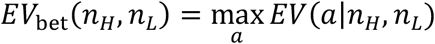

where *α* ∈ {*α_H_*, *α_L_*} and

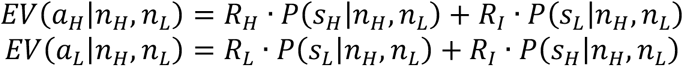

Since the posterior is determined by the beads difference (Eq. 1), the bet *EV*s are also determined by the beads difference.

In order to evaluate *EV*_draw_, we have to take into account two important facets of our information-seeking paradigm: first, the content of information (the type of the next bead, *b_H_* or *b_L_*) is stochastic, and second, the agent can decide whether to draw yet another bead or not after observing the next bead. Therefore, we have to evaluate the likelihood of the next bead type and combine it with the *EV* of an optimal choice conditional on each bead type. The likelihood of the next bead type based on the current evidence is evaluated according to the posterior on the jar type:

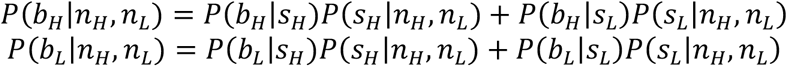

If the next bead is *b_H_*, it would update the evidence from (*n_H_*, *n_L_*) to (*n_H_* + 1, *n_L_*). Then the agent can either make an optimal bet and achieve *EV*_bet_(*n_H_* + 1, *n_L_*) or pay the cost to draw another bead and achieve *EV*_draw_(*n_H_* + 1, *n_L_*) − *c*_draw_. Similarly, if the next bead is *b_L_*, it would update the evidence to (*n_H_*, *n_L_* + 1), based on which the agent can either make an optimal bet and achieve *EV*_bet_(*n_H_*, *n_L_* + 1) or draw another bead and achieve *EV*_draw_(*n_H_*, *n_L_* + 1) − *c*_draw_. Therefore, the highest *EV* that the agent can achieve after drawing an additional bead is

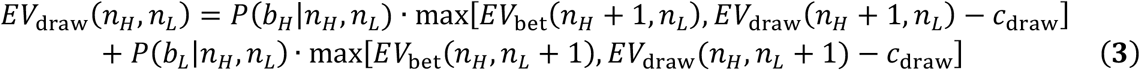

In Eq. 3, *EV*_draw_(*n_H_*, *n_L_*) in the left-hand side depends on *EV*_draw_(*n_H_* + 1, *n_L_*) and *EV*_draw_(*n_H_*, *n_L_* + 1) in the right-hand side due to the aforementioned sequentiality of information seeking. We thus solved Eq. 3 by backward recursion. Specifically, we arbitrarily assumed that the agent cannot draw more than 200 beads, set *EV*_draw_(*n_H_*, *n_L_*) = 0 where *n_H_* + *n_L_* = 200, and used Eq. 3 to obtain *EV*_draw_(*n_H_*, *n_L_*) where *n_H_* + *n_L_* = 199. We then used Eq. 3 recursively to obtain *EV*_draw_(*n_H_*, *n_L_*) for all cases where 0 < *n_H_* + *n_L_* < 200. Although the obtained *EV*_draw_(*n_H_*, *n_L_*) depends on *n_H_* + *n_L_*, it reaches an asymptote over the course of recursion quickly (Fig. S2). We substituted the asymptotic *EV*_draw_ to Eq. 2 and obtained the theoretical *VOI* as a function of the beads difference.

The *VOI* obtained for each of the three reward structures, (*R_H_*, *R_L_*, *R_I_*) = (70, 10, 0), (170, 110, 100), and (7, 1, 0), is shown in Fig. 2F. The baseline shift affects both *EV*_draw_ and *EV*_bet_ by the same amount, which is canceled out in Eq. 2 and does not affect the *VOI*. On the other hand, since *c*_draw_ was not scaled along with rewards and remained the same across conditions (0.1 points), the scale manipulation affects not only the magnitude but also shape of *EV*_draw_ (Eq. 3) and thus the *VOI* (Eq. 2).

The most important prediction of this theoretical framework is that *information seeking should be biased due to the reward asymmetry*. The *VOI* takes an inverted-*U* shape as a function of the beads difference, and its peak is at a moderate negative beads difference (*n_H_* − *n_L_* = −5). This is because the information would directly improve the subsequent bet choice; when *n_H_* − *n_L_* = −5, *EV*(*α_H_*|*n_H_*, *n_L_*) is close to *EV*(*α_L_*|*n_H_*, *n_L_*), but the next bead would increase their difference in either direction (if a high-reward bead *b_H_* is observed, *EV*(*α_H_*|*n_H_* + 1, *n_L_*) > *EV*(*α_L_*| *n_H_* + 1, *n_L_*); if a low-reward bead *b_L_* is observed, *EV*(*α_H_* | *n_H_*, *n_L_* + 1) < *EV*(*α_L_* | *n_H_*, *n_L_* + 1)). Therefore, the agent can bet on *S_H_* after *b_H_* and bet on *S_L_* after *b_L_*, and such flexibility improves the overall *EV*. In contrast, the *VOI* is effectively zero when the beads difference is positive (*n_H_* − *n_L_* > 0), because the agent would bet on *S_H_* irrespective of the next draw. The *VOI* is also effectively zero when low-reward beads outnumber high-reward beads by a large enough margin (*n_H_* − *n_L_* < −7), because the agent would bet on *S_L_* irrespective of the next draw.

This qualitative prediction, a bias in information seeking towards a negative beads difference, does not depend on most of our assumptions (e.g., choice optimality, risk neutrality). Information seeking would be biased *as far as the agent is sensitive to the rank order of rewards and the bead difference*. On the other hand, if an agent is not motivated to maximize rewards but to maximize the accuracy of the prediction (i.e., their utility function *U* follows *U*(*R_H_*) = *U*(*R_L_*) > *U*(*R_I_*)), they would exhibit unbiased information seeking; the uncertainty about the jar type is determined by |*n_H_* − *n_L_*| and is highest when *n_H_* = *n_L_*, which is when the agent would draw beads most frequently. Therefore, a bias in information seeking would suggest that information seeking is motivated by information’s instrumentality for future reward seeking.

### Behavioral data analysis

In order to examine information-seeking behavior, we analyzed the frequency at which participants drew at least one bead as a function of the beads difference. We specifically focused on whether they drew the first bead as a function of the current evidence and examined if it was biased by the reward asymmetry as theoretically predicted. The relationship between information-seeking behavior and the beads difference was analyzed using Gaussian Process (GP) logistic regression (Rasmussen & Williams, 2006). GP logistic regression estimates a latent function that smoothly varies with the independent variable (the beads difference) and yields likelihoods of binary choices (whether participants drew a bead in each trial), and the estimated latent function can be interpreted as the subjective *VOI* function (the higher *VOI* is, the more likely participants draw a bead). The latent function with isotropic squared exponential covariance was estimated using Variational Bayes approximation, as implemented in Gaussian Processes for Machine Learning toolbox, version 4.2 (https://github.com/alshedivat/gpml) (Rasmussen & Nickisch, 2010).

To test whether information-seeking behavior systematically differed across blocks and reward conditions within each block, we compared four models. Model 1 implemented the theoretical prescription that information seeking is sensitive to the scale manipulation but not to the baseline manipulation. It thus consisted of three separate latent value functions, one used in all trials in the baseline block, one used in trials where (*R_H_*, *R_L_*, *R_I_*) = (70, 10, 0) in the scale block, and one used in trials where (*R_H_*, *R_L_*, *R_I_*) = (7, 1, 0) in the scale block. We constructed several alternative models. Model 2 postulated different value functions for reward conditions not only in the scale block but also in the baseline block, one for trials where (*R_H_*, *R_L_*, *R_I_*) = (70, 10, 0) and another for trials where (*R_H_*, *R_L_*, *R_I_*) = (170, 110, 0) (i.e., four value functions in total); Model 3 postulated the lack of sensitivity to reward conditions in both blocks but a separate value function for each block (i.e., two value functions in total); and Model 4 postulated one common value function for all trials in both blocks. These models were compared based on leave-one-participant-out cross validation (LOPO CV) and leave-one-trial-out cross validation (LOTO CV). We also adopted the same analytic approach to the bet choices, comparing the performance of Models 1-4.

We found that Model 3 outperformed other models for both information-seeking and bet choices (see Results). To test whether information-seeking behavior was biased by the reward asymmetry, we next compared Model 3 with another model (Model 5) that assumed value functions that are symmetric with respect to the beads difference (i.e., value functions that only vary with the absolute value of beads difference). We found that Model 3 fit information-seeking behavior better than Model 5, supporting a bias in information seeking (see Results).

The fact that Model 3 performed better than Models 1, 2, and 4 suggests that, while participants did not change their behavioral strategies based on the trial-by-trial reward manipulation, they adapted to the different reward statistics across blocks. However, such changes across blocks could potentially reflect time-induced behavioral changes as well, such as boredom or fatigue, since all participants completed the baseline block first and the scale block second. To examine the possibility that the population-level behavioral pattern was not stationary over time, we tested another model (Model 6) that assumed distinct value functions between the first and second scanning runs within each block (one value function for each run, four functions in total). Model 6 performed worse than Model 3 (information-seeking choices: LOPO CV log likelihood [LL] = −1222.05 vs. −1214.73, LOTO CV LL = −1145.39 vs. −1142.25, bet choices: LOPO CV = −288.55 vs. −283.56, LOTO CV LL = −266.00 vs. −265.26), suggesting that changes in participants’ behavior were systematically driven by reward statistics rather than time.

### MRI data acquisition

MRI data was collected using a Siemens (Erlangen, Germany) Trio 3T scanner with a 32-channel head coil at the University of Pennsylvania. A 3D high-resolution anatomical image was acquired using a T1-weighted MPRAGE sequence (voxel size = 0.9375 × 0.9375 × 1 mm, matrix size = 192 × 256, 160 axial slices, TI = 1100 msec, TR = 1810 msec, TE = 3.51 msec, flip angle = 9 degrees). Functional images were acquired using a T2*-weighted multiband gradient echoplanar imaging (EPI) sequence (voxel size = 2 × 2 × 2 mm, matrix size = 98 × 98, 72 axial slices with no interslice gap, 400 volumes, TR = 1500 msec, TE = 30 msec, flip angle = 45 degrees, multiband factor = 4), followed by Fieldmap images (TR = 1270 msec, TEs = 5 msec and 7.46 msec, flip angle = 60 degrees).

### MRI data analysis

MRI data were analyzed using FSL (FMRIB Software Library, version 6.0) (Jenkinson et al., 2012; Smith et al., 2004). MPRAGE anatomical images were skull-stripped using FSL BET. EPI functional images were slice-time corrected, motion corrected (FSL MCFLIRT), high-pass filtered (cutoff = 90 sec), geometrically undistorted using Fieldmap images, registered to the MPRAGE anatomical image, normalized to the MNI space, and spatially smoothed (Gaussian kernel FWHM = 6 mm).

To look for regions that represent the subjective *VOI* upon the initial beads presentation, we ran a GLM analysis (GLM 1). The regressor of interest modeled the initial beads presentation (3-second boxcar) and was parametrically modulated by the trial-by-trial subjective *VOI*, which was the latent function estimated in the winning model (Model 3) of GP logistic regression on the information-seeking behavior. GLM 1 also included nuisance regressors that modeled the initial beads presentation (unmodulated), the extra bead presentation, and button presses. The regressors were convolved with the canonical double-gamma hemodynamic response function (HRF). GLM 1 additionally incorporated six head motion parameters (3 translations and 3 rotations, estimated by MCFLIRT) as confound regressors. GLM 1 was run following the standard approach of FSL FEAT; the GLM was first fit to BOLD signals in each run (first level) and the estimated coefficients of interest were combined across runs (second level). Individual-level *T*-statistics were entered into the population-level inference using FSL randomise, in which clusters that showed positive response to subjective *VOI* were defined at the voxel-wise cluster-forming threshold of *p* < .001 and evaluated by sign-flipping permutation on cluster mass. A cluster that survived whole-brain family-wise error (FWE) corrected *p* < .05 is reported in Fig. 3B; another cluster that survived a more lenient threshold (*p* < .10) is reported in Fig. S1.

To illustrate how the cluster’s activation varied as a function of the beads difference, we ran another GLM (GLM 2) using FSL FEAT, which included a regressor for each level of beads difference separately, along with the same nuisance regressors as GLM 1. *T*-statistics for each regressor of interest were then averaged across runs within each block and then averaged across all voxels in the right DLPFC cluster defined as above (Fig. 3B).

Lastly, to examine how the DLPFC responds to the updating of *VOI*, we ran another GLM (GLM 3) using FSL FEAT to estimate the time course of signals related to the initial *VOI* and the *VOI* updating, which were derived from Model 3 of GP logistic regression. The *VOI* updating was calculated as the signed difference between the posterior *VOI*, which depends both on the initial beads and the extra bead, and the prior *VOI*, which depends only on the initial beads (Fig. 4A). GLM 3 included three sets of finite impulse response (FIR) function, one unmodulated (intercept), one parametrically modulated by the initial *VOI*, and one parametrically modulated by the *VOI* updating. These FIRs were aligned to the onset of the extra bead and sampled every 1.5 seconds (equal to TR) for the total duration of 21 seconds. GLM 3 also included nuisance regressors that modeled the initial beads presentation and button presses, convolved with the canonical HRF, along with head motion parameters. *T*-statistics of parametrically modulated FIR sets were averaged across all voxels in the right DLPFC cluster for each participant. Population-level inference on the updating signal was conducted at the cluster level across time; clusters were defined at the event-wise cluster-forming threshold of *p* < .05 and evaluated by sign-flipping permutation on cluster mass, correcting for FWE across time.

## Supplementary Figures

**Fig. S1.**
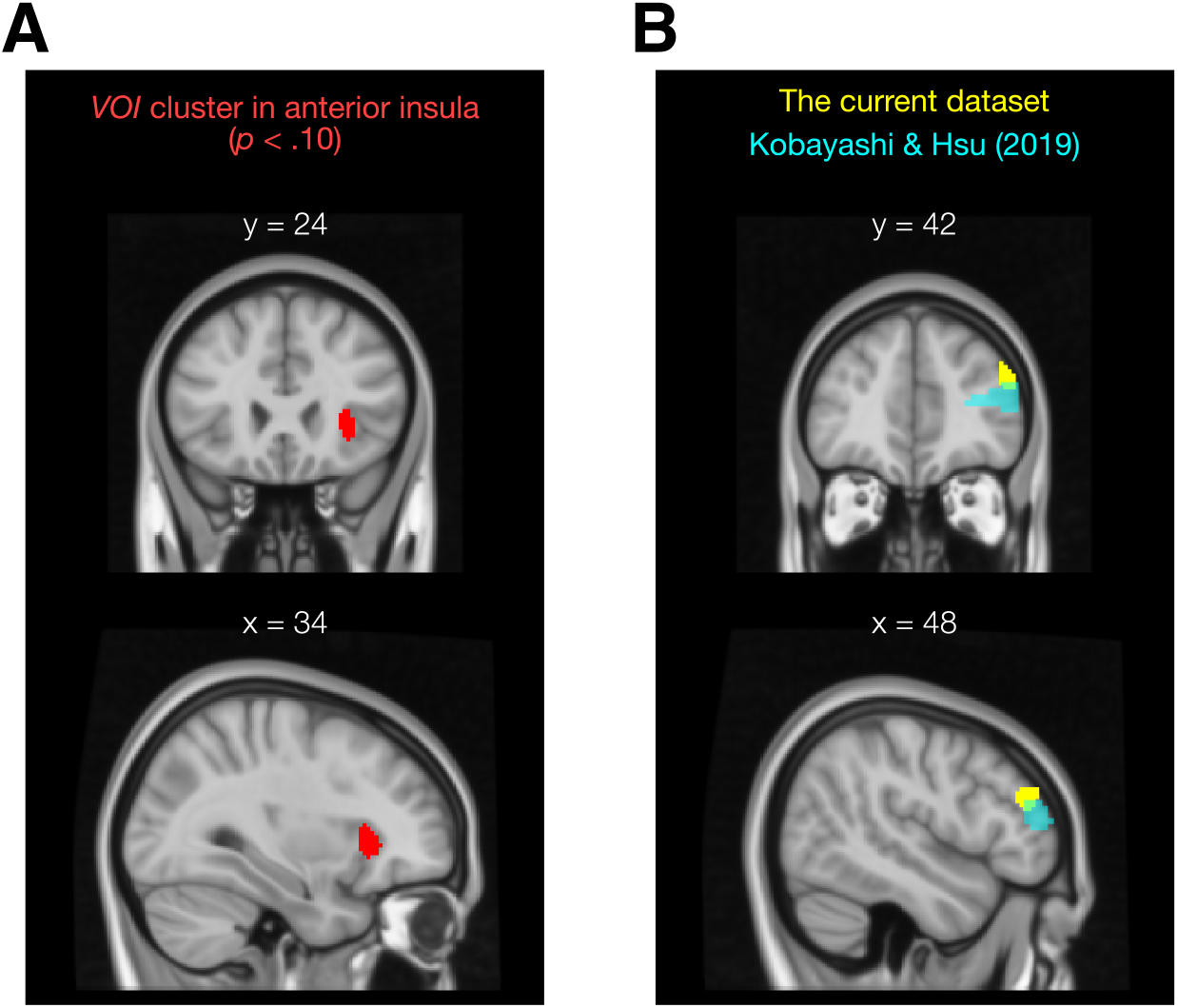
(A) At a liberal threshold (cluster-forming threshold *p* < .001, cluster mass *p* < .10, corrected for whole-brain FWE), the subjective *VOI* was positively associated with activations in right anterior insula. (B) The DLPFC cluster identified in the current dataset (Fig. 4b) overlaps with a subjective *VOI* cluster reported in Kobayashi & Hsu (2019).

**Fig. S2.**
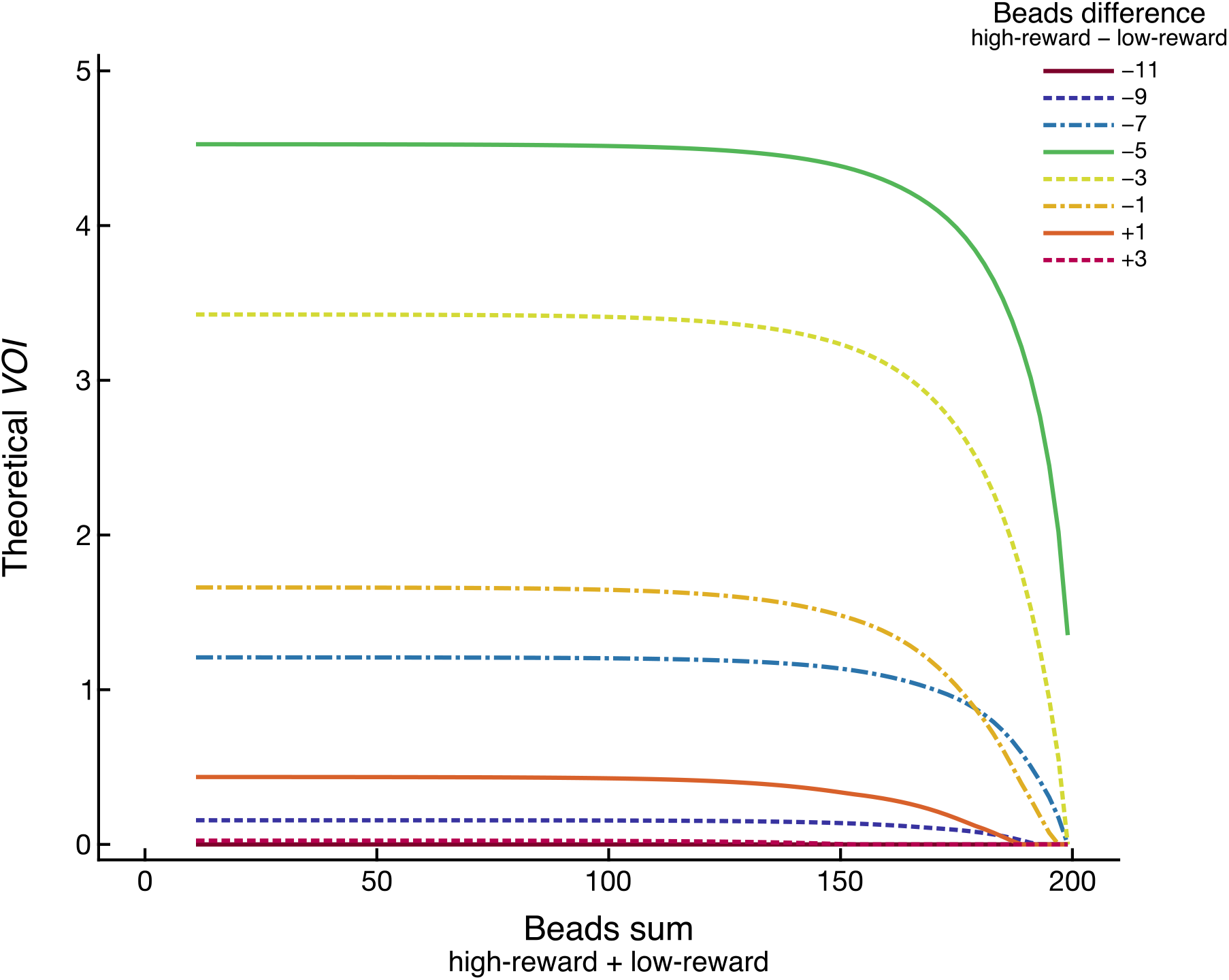
The theoretical *VOI* was numerically estimated by backward recursion (up to 200 steps). The *VOI* reached an asymptote at each level of beads difference over the course of recursion. Moreover, the *VOI* was highest with a negative beads difference (−5) throughout recursion.

